# Direct observation of rotation-coupled protein diffusion along DNA on the microsecond timescale

**DOI:** 10.1101/401414

**Authors:** Emil Marklund, Elias Amselem, Kalle Kipper, Xuan Zheng, Magnus Johansson, Sebastian Deindl, Johan Elf

## Abstract

Many proteins that bind specific DNA sequences search the genome by combining three dimensional (3D) diffusion in the cytoplasm with one dimensional (1D) sliding on non-specific regions of the DNA^1–5^. It is however not known how sliding proteins are oriented with respect to DNA in order to recognize specific sequences. Here we measure the polarization of fluorescence emission from single fluorescently labeled *lac* repressor (LacI) molecules sliding on stretched DNA. Real-time feedback-coupled confocal single-particle tracking allows us to measure fluorescence correlation of the sliding molecules. We find that the fluctuations in the fluorescence signal on the μs timescale are accurately described by rotation-coupled sliding on DNA. On average, LacI moves ∼50 base pairs per revolution, which is significantly longer than the 10.5 bp helical periodicity of DNA. Our data support a facilitated diffusion model^1^ where the transcription factor (TF) scans the DNA grooves for hydrogen bonding opportunities in a pre-aligned orientation with occasional slippage out of the groove.

---

Not much is known about the dynamic nature of protein-DNA interactions during sliding. Ensemble methods such as x-ray crystallography^6^ and NMR spectroscopy^7^ have provided important insights into the structure and orientation of proteins interacting with specific DNA sequences. However, the orientational properties of proteins sliding on DNA have only been inferred by indirect methods. For example, the diffusion rate constants of some sliding proteins scale with their sizes in a manner consistent with translocation coupled rotation around the DNA^8^. However, this model relies on the assumptions that the different proteins all have approximately the same interaction energy with DNA and slide along it with the same helical pitch. There are also examples of proteins whose translocation on DNA appears inconsistent with a rotation-coupled model^9^. Here we have used single-molecule-tracking aided fluorescence correlation spectroscopy to measure the *lac* repressor’s mode of sliding and pitch of rotation during translocation on flow stretched DNA^3,10^.

As a readout for protein orientation during LacI sliding we used the dipole field emitted by a linked dye. To reduce rotation of the dye in relation to the protein, we covalently bound bifunctional rhodamine B (BR)^11^ to two engineered cysteine residues in the DNA binding domain of LacI (Figure 1a and Extend Data Table 1). Molecular dynamics simulations of the DNA-LacI-BR complex showed that that the bifunctional labeling is more rigid compared to monofunctional labeling with a Cy3 attached at one cysteine (Figure 1b). However, both dyes have dipole moment distributions that are far from uniform making them well-suited candidates for measuring ortientational properties. The labeled LacI dimer (Extended Data Fig. 1) retained the ability to bind the *lac* operator (Extended Data Fig. 2). To monitor sliding events (Extended Data Table 2-3) on flow-stretched DNA we anchored one end of the 49 kB λ-DNA to a poly(ethylene glycol) (PEG)-passivated coverslip^12^ *via* a biotin-neutravidin linkage and assembled a fluidic channel (Figure 1c).

**Figure 1.**
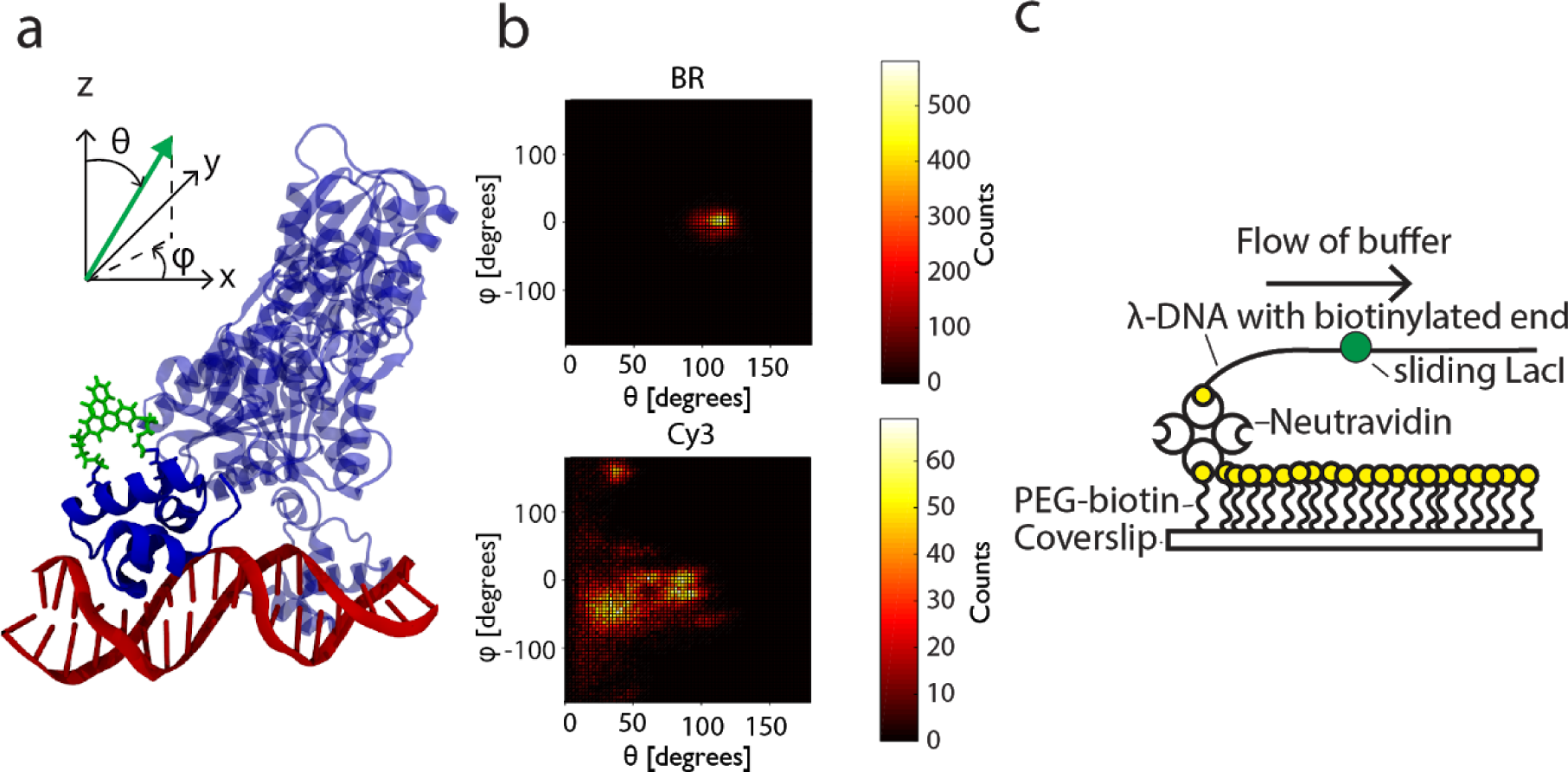
Bifunctionally labeled LacI on flow stretched DNA. a. Molecular model of the studied system. Bifunctional rhodamine (green) attached to an α-helix on the DNA (red)-bound *lac* repressor (blue) via two cysteine residues. b. Spherical coordinate (polar angle *θ* and azimuthal angle *φ*) distributions showing rigidity of BR (top) and Cy3 (bottom) in 70 ns molecular dynamic simulations. c. Schematic for the DNA flow-stretching experiment in a fluidic channel.

**Figure 2.**
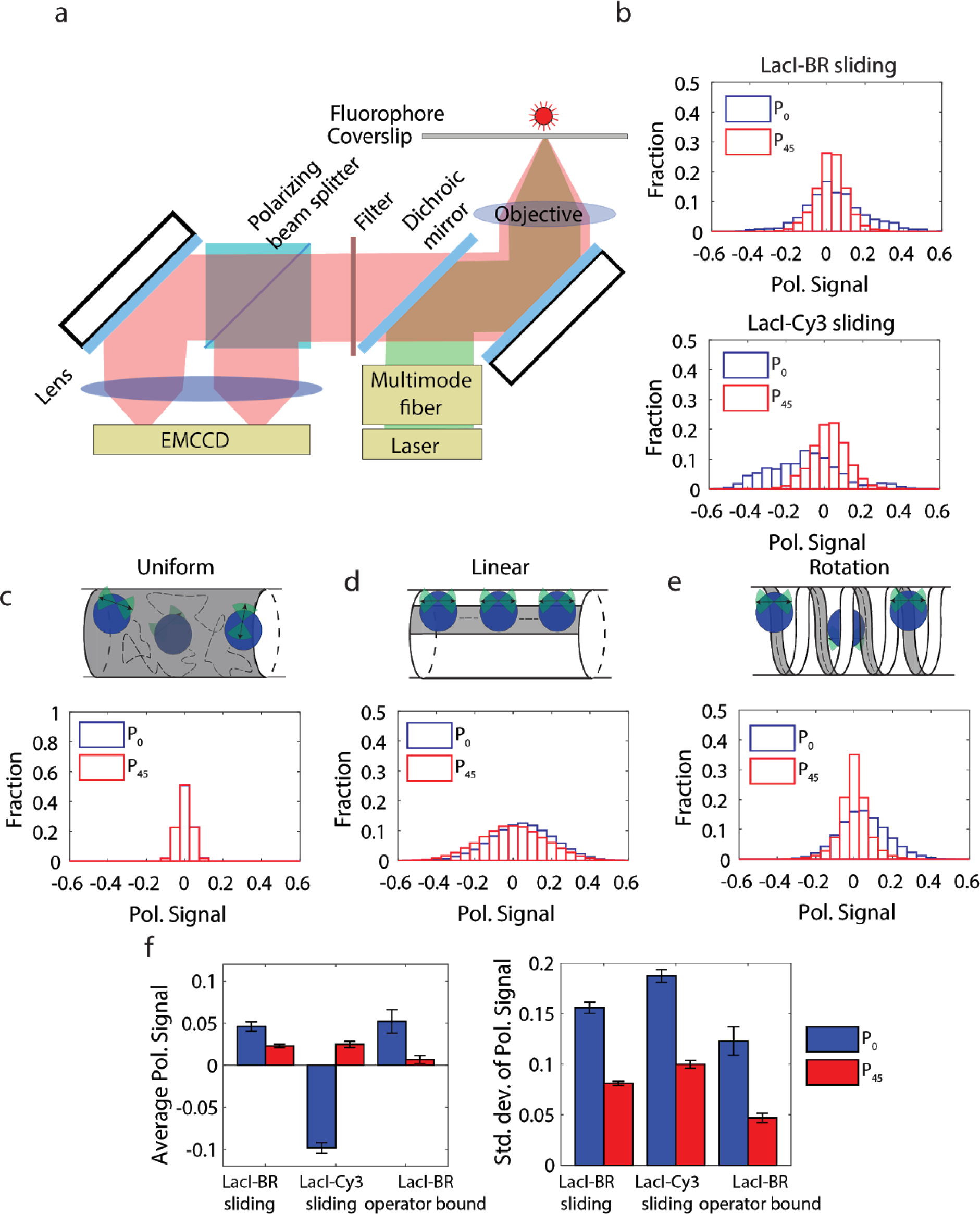
Polarization measurements. a. Schematic illustration of the experimental setup. b. *P*_*0*_ (blue) and *P*_*45*_ (red) polarization distributions averaged over 600 ms (3 frames) per count measured for sliding LacI-BR (top) and LacI-Cy3 (bottom). 79 and 172 (LacI-BR), respectively 61 and 53 (LacI-Cy3) sliding events were detected in the *P*_*0*_ and *P*_*45*_ measurements, respectively. c-e. Simulated polarization distributions for the uniform (c), linear (d), and rotation-coupled (e) sliding models. f. Averages (left) and standard deviations (right) of experimental polarization distributions. Error bars represent standard errors.

First we investigated the orientational properties of sliding LacI using single-molecule fluorescence with widefield epi-fluorescence and camera-based polarization detection^13–17^ (Figure 2a). In these measurements the observed polarization signal is an average determined by all orientations that the protein adopts while sliding over hundreds of basepairs during the acquisition time of 200 ms. The polarization measurement of the fluorescence emission is detected relative to the DNA stretching direction (α). Both polarization components *I*_*α*_ and *I*_*α+90*_ are detected simultaneously on an EMCCD and used to calculate the polarization signal as

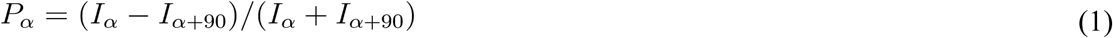

The polarization signal for individual LacI molecules sliding on DNA was measured with one polarization axis aligned with the stretching direction of DNA (measuring *P*_*0*_), as well as shifted 45° from it (measuring *P*_*4*5_) (Figure 2b). These camera-based polarization measurements can be used to rule out some modes of sliding. The rationale for the analysis is that the polarization signal is averaged out and thus close to zero in the directions corresponding to axes of symmetry where the protein is free to rotate. On the other hand a preferential orientation would manifest itself in a broadening of the distribution of polarizations. For example, if the protein was free to move and rotate on the DNA surface in all directions at a time scale faster than the imaging frame rate, the polarization distribution would be narrow in both *P*_*0*_ and *P*_*45*_. As a reference, this case was simulated using a vector-based point-spread function model for translating dipole orientation^18^ (Figure 2c). In contrast, if the protein did not rotate around any of its own axes and only diffused linearly along the DNA any bias in direction of the DNA, protein or dye on the imaging time-scale would give rise to non-zero polarization signals resulting in a broadening in both *P*_*0*_ and *P*_*45*_ (Fig. 2d). Furthermore, *P*_*0*_ would have a non-zero average if the dye exhibited a preferred orientation with respect to DNA whereas *P*_*45*_ would be indifferent with respect to DNA-protein alignment. The addition of a fast rotation of the protein around the DNA axis would not change *P*_*0*_ compared to the linear diffusional sliding, whereas *P*_*45*_ would be averaged out due to the symmetry of the rotation (Fig. 2e). The experimental distributions for LacI labeled with rhodamine or Cy3 (LacI-BR and LacI-Cy3) both display broadening in *P*_*0*_ as compared with *P*_*45*_. When comparing with the simulations, this experimental result is only compatible with rotational diffusion around the DNA axis. We also note that the *P*_*0*_ distribution is positive for LacI-BR and negative for LacI-Cy3. This means that BR is more aligned with the DNA than Cy3, which agrees with our molecular dynamics simulations.

To investigate the sources of noise in the measurement, we compared the polarization of the sliding LacI-BR to that of operator bound LacI-BR (Fig. 2f and Extended Data Fig. 3). We found that their average *P*_*0*_ polarization signals were very similar, suggesting that the protein may slide in an orientation that is close to the operator bound structure. The *P*_*0*_ standard deviation was slightly smaller for the operator bound conformation, showing that only some of the broadening could be attributed to protein DNA interactions while sliding. Most importantly however, *P*_45_ was averaged out also for the operator bound molecule suggesting that DNA itself was rotating around its own long axis on the imaging time scale.

**Figure 3.**
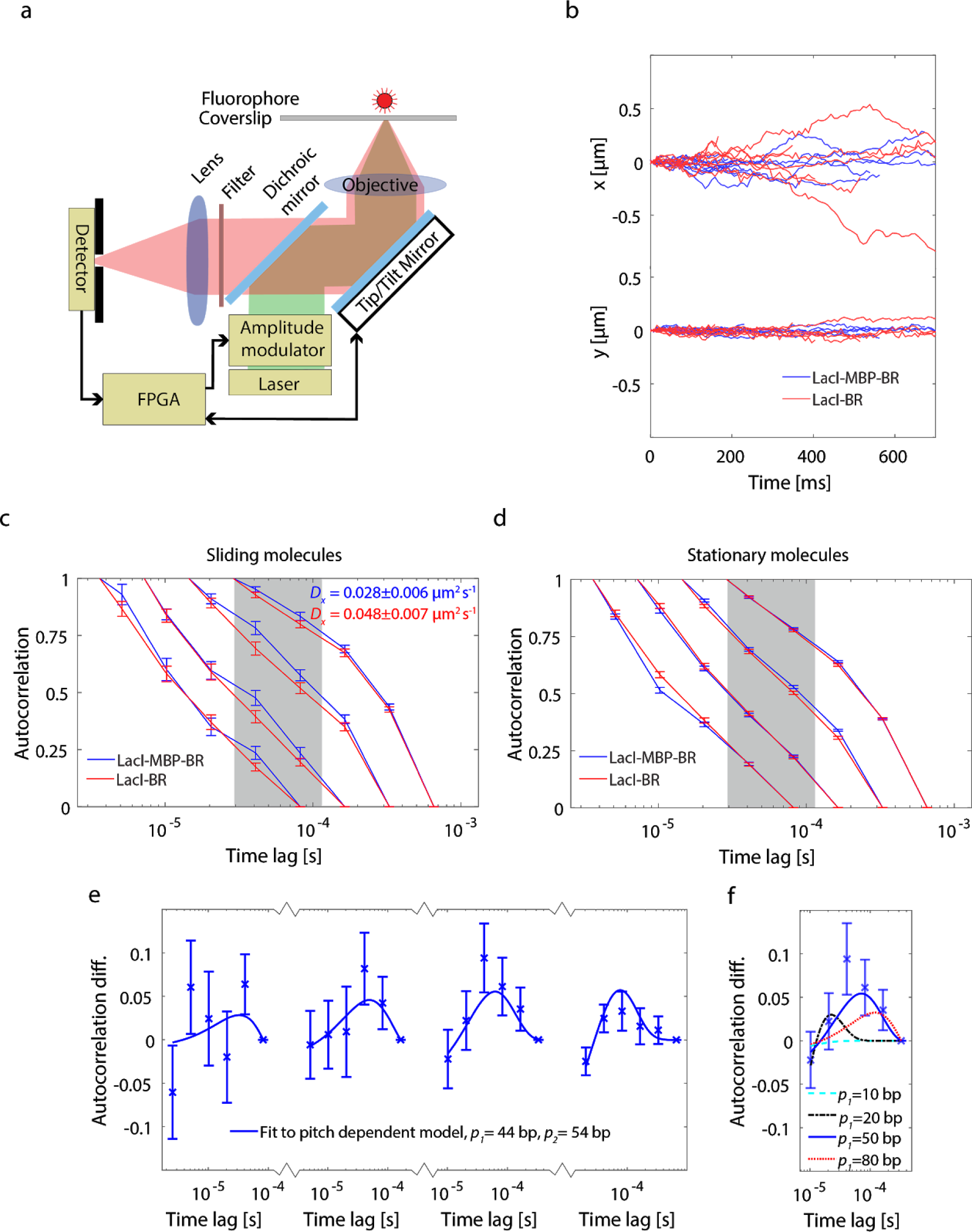
Single-molecule-tracking aided fluorescence correlation spectroscopy. a. Schematic illustration of the experimental setup. b. x (DNA direction) and y coordinate of sliding LacI-MBP-BR and LacI-BR molecules obtained by the confocal tracking system. In total, 93 and 118 sliding molecules were captured for LacI-MBP-BR and LacI-BR respectively. For clarity, 10 representative traces are shown for each species. The average normalized autocorrelation of the fluorescence signal for sliding (c) and stationary (d) LacI-MBP-BR and LacI-BR molecules. The reported diffusion constants are averages of all tracked molecules. Error bars and error estimates represent standard errors. e. The difference in autocorrelation between LacI-MBP-BR and LacI-BR in the four different time regimes. Error bars represent standard errors. Lines represent the best fit to a rotation-coupled sliding model. The fit yield pitches for the rotation of 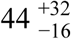 (LacI-MBP-BR) and 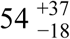 (LacI-BR) base pairs with 95 % confidence intervals. f. Best fits of the rotation-coupled sliding model to the experimental data in the third time regime when the pitch of LacI-MBP-BR is constrained at four different levels.

The temporal resolution in camera-based tracking limited us to study orientational properties averaged over hundreds of basepairs of sliding, which made it impossible to distinguish between a rotation of the DNA around its own long axis and a rotation of LacI around the DNA. To study the protein rotation around DNA, we therefore tracked the sliding LacI-BR molecules (Extended Data Table 4-5) in real-time with a focused laser (Figure 3a and Extended Data Fig. 4) while we at the same time recording the fluctuations in the fluorescence signal on the microsecond time scale using fluorescence correlation spectroscopy (FCS)^19^ (Extended Data Fig. 5). This enabled us to measure both the translational (Fig. 3b) and rotational diffusion rates.

Each detected photon was time-tagged on the nanosecond time scale and used to determine the autocorrelation in the fluorescence signal in several different time regimes starting at 3 µs (Figure 3c). We reasoned that if LacI sliding were coupled to its rotation around the DNA, the fluctuations in fluorophore orientation should be correlated with the rate of translational diffusion. A decreased diffusion coefficient is therefore expected to increase the decay time in the autocorrelation in the time regime relevant for rotation-coupled sliding. Indeed, when the experiment was repeated with a larger variant of LacI (LacI fused with maltose binding protein) labeled with the same fluorophore (LacI-MBP-BR), we measured slower translational diffusion as well as slower decay in the autocorrelation of the fluorescence signal in the 30 to 100 µs range (Figure 3c). This result depended on sliding as the autocorrelation of the fluorescence signals for stationary molecules stuck on the glass surface did not exhibit these differences between the two LacI species (Fig 3d). Our data thus show that LacI sliding is coupled to its rotation around the DNA, with characteristic decay times in the order of 40 µs where the largest autocorrelation difference between the two LacI variants is observed.

To estimate the distance that LacI translocates along the DNA during one full rotation, we fit the difference in autocorrelation from all time-scales to a model where the pitches of the helical rotation of the two proteins are free parameters (Figure 3e-f). The fitting method accurately returned the correct pitches when it was tested on simulated data where the pitches were known and the background was taken from the autocorrelation of stationary LacI molecules (Extended Data Fig. 6). When applied to the experimental data, fitting yielded estimates for the pitch of 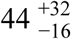 bp and 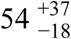 bp for LacI-BR and LacI-MBP-BR respectively.

The structural dynamics of protein DNA interactions on the microsecond time scale shows that the lac repressor rotates around the DNA helix while sliding with a ∼50 bp pitch, which is longer than the 10.5 bp periodicity of the DNA. A model that explains these observations combines faithful protein tracking of the DNA helix with occasional inter groove slippage. Such slippage may be a requirement for keeping non-specific binding sufficiently weak to optimize overall search speed^1^ and result in the previously suggested bypassing of specific binding sites^2^. The technology developed for this study can be extended to tracking molecules diffusing and interacting inside cells, so that classification of binding states that are currently determined by rates of translational diffusion only, can be complemented with spectroscopic information such as the rotational diffusion profile.

## Methods

### Protein purification, labeling and functionality testing

All LacI constructs (Extended Data Table 1) were designed with a C-terminal 6xHis-tag for purification purposes and a truncated tetramerization domain with a design that previously has been shown to retain the ability of the protein to form dimers^20^. Additional cysteines were introduced into the amino acid sequence to enable monofunctional (LacI-Cy3) or bifunctional (LacI-BR and LacI-MBP-BR) labeling. Detailed descriptions on the purification and labeling has previously been published for LacI-Cy3^20^ and can be found in the Supplementary Information for LacI-BR and LacI-MBP-BR. In addition to the single molecule experiments performed in this study, functionality was tested by measuring the affinity to the O1 operator for LacI-BR (Extend Data Table 1) in a gel-shift assay (see Supplementary Information for details).

### DNA flow stretching and single-molecule microscopy sample preparation

Preparation of flow channels and tethering and stretching of dsDNA by laminar flow was carried out according to previously published methods^12,21^. The 49 kB λ-DNA was used for sliding experiments and a 7kb DNA with an Osym operator in the middle and Cy3 fluorophores in each end (created by PCR) was used for operator binding experiments. Sliding experiments were performed with a fluorescent protein concentration between 10 and 100 pM. Operator binding experiments were performed by pre-incubating the Osym DNA and LacI at a high concentration, followed by dilution and introduction of the complex into the flow-channel at a fluorescent protein concentration of 30 pM (see Supplementary Information for details).

### Single-molecule fluorescence polarization microscopy

Schematics of the optical setup are shown in Figure 2a of the main-text and in more detail in Extended Data Figure 4. A supercontinuum laser (NKT EXW-4) was used as excitation laser. The optical spectra was filtered down to 540±18 nm by the filter combination Semrock FF01-770SP-25 and Semrock FF01-532-18. The light was brought to the custom built microscope through a multimode fibre (Thorlabs M69L05) with a mode scrambler. The mode scrambler both randomizes the polarization and diffuses the light source. Sample excitation was done through an EPI-illumination configuration using a dichroic mirror (Semrock FF552-Di02) and a 100x Nikon objective (CFI Plan Apo Lambda, NA=1.45). Fluorescence was collected through a dichroic mirror (Semrock FF552-Di02) and filter (Semrock FF01-585-40) which was followed by a 2x expander (Thorlabs achromatic doublets f = 100 mm and f = 200 mm) to obtain a final total magnification of 200x and a pixel size of 80 nm. The emitted light was divided into its horizontal and vertical components by two relay lenses (Thorlabs achromatic doublets f = 150 mm). Each polarization component was imaged on separate part of a EMCCD camera (Princeton Instruments PhotonMAX 512). The laser intensity over the sample was set to 3 kW/cm^2^ and the EMCCD was configured to have an exposure time of 160 ms which resulted in a 5 fps frame rate. The EMCCD gain was optimized to obtain an average signal to noise ratio of 1.6.

### Single-molecule-tracking aided fluorescence correlation spectroscopy

#### Optical setup

Schematics of the optical setup are shown in Figure 3a of the main-text and in more detail in Extended Data Figure 4. After the spectral filtration the supercontinuum laser light was brought to an amplitude modulator, constituting of two polarizers with a pockel cell (Newport 4102NF) in between. A spatial filter (11mm lens, 10 μm pinhole followed by a 75 mm lens) was used to obtain a well behaving Gaussian beam. A tip tilt piezo mirror (Piezosystem Jena PSH 10/2) was placed after the dichroic mirror at the back focal plane of the objective. The excitation laser was set to circular polarization in the xy-plane by rotating a combination of a half and quarter wave plates and measuring the polarization after the objective. The intensity at the back focal plane of the objective was set to 15μW and emitted fluorescence was collected through the dichroic mirror followed by an imaging lens, a 75 μm pinhole, polarizing beam splitter (PBS) and an avalanche photodiode (APD, SPCM-AQRH Excelitas) for each polarization. A field programmable gate array (FPGA, NI 7852R) was used for sampling each APD signal, time tagging photon arrival times, triggering laser excitation pulses and controling the real-time feed-back of the piezo tip/tilt mirror.

#### Single-molecule-tracking and photon counting data acquisition

The tracking system was started as a raster scan. The scanning works both as a standard confocal imaging system and a search for potential tracking events. Here the excitation laser was triggered with a 500 μs pulse with a sampling rate of 1900 samples/s. If the photon count was higher than a set threshold the system entered the tracking mode. In the tracking mode the tip/tilt piezo mirror traced out a circle centered around the position where the photon count passed the threshold requirement with a revolution time of 4 ms and a diameter of 700 nm. During each loop the laser is triggered 6 times over equally spaced intervals each with a duration of 500 μs. The fluorophore offset from current estimated positions was triangulated by calculating a rolling mean weighted centroid using the 12 latest points. A PID controller took the offset value and generated a correction for the current position followed by the triggering of a position update of the piezo tip/tilt mirror with the new corrected fluorophore position. This was performed for each new measurement point and the tracking scheme was repeated until the mean photon count dropped under a set threshold, terminating the tracking loop and bringing the system back into the raster scanning mode. Details about the optimization of tracking parameters can be found in the Supplementary information.

Simultaneously, at the beginning of each 500 μs long laser pulse a 200 Mhz counter was started. Each time a photon was detected by an APD the counter value together with the APD id was registered and saved. With this time-tagged time-resolved (TTTR) data it is possible to reconstruct the photon time trace over a tracking trajectory with a time resolution of 5 ns and perform FCS analysis.

### Data analysis

#### EMCCD polarization microscopy data analysis

The two polarization components emitted from the same fluorophore were first correlated to each other by translational registration of a 10 μm x 10 μm grid, imaged prior to each experiment. Detection and localization of single-molecules were then done on the sum of the two polarization components. An à trous wavelet three-plane decomposition was used for dot detection^22^. The dots were detected through scale-dependent standard deviation-thresholding in the third wavelet plane, where the standard deviation was estimated by the median absolute deviation method^23^ and the threshold was set to 3 standard deviations. Dot centers were localized by calculating the weighted centroid from the pixel regions obtained from the dot detection. Centroids in different frames of the movie were then connected to trajectories with u-track^24^ using a maximal search radius of 4 pixels for the sliding experiments and 3 pixels for the operator binding experiments. To account for fluorophore blinking, a maximal dark time of 2 frames were used in the sliding experiment, while no upper limit was set on the dark time in the operator binding experiments (trajectory building in the operator experiments is more trivial since the dots do not move)

Classification of sliding trajectories were done with principal component analysis (PCA) on trajectories longer than 15 frames (3 s). Trajectories were only classified as sliding if their principle direction of movement did not deviate more than 20° from the expected DNA stretching direction, if the eigenvalue of largest principal component was larger than 0.25 μm^2^s^−1^, and if the eigenvalue of the smallest principal component was smaller than 0.06 μm^2^s^−1^ (see Extend Data Table 3 for more classification parameters). The classifier was optimized to effectively find sliders in flow-channels with DNA but not in negative control channels without DNA (Extended Data Table 2).

Operator binding events were analyzed by selecting the trajectories corresponding to the LacI dot from the EMMCD data (located approximately 4 kb ∼ 13 pixels and 3 kb ∼ 10 pixels away in the DNA stretching direction from the 5’- and 3’- Cy3 dot respectively).

The background intensities of pixels were estimated by calculating a moving average of each fluorescence image, implemented with exclusion of outliers to not include fluorescent dots in the average. The two polarization intensities of single-molecules in each frame were calculated as the difference between the raw fluorescence intensity and the background, summed over a 7 × 7 pixel square around the centroid of the dot. The polarization signal was then calculated according to equation (1) after summation of the intensities over 3 frames (600 ms).

#### Trajectory building of confocal tracking data

The outputs available from the real-time tracking are the tip/tilt mirror position for each 500 μs laser pulse, the total photon count during the same laser pulse interval and a flag showing if the tracking mode was active. In the parts of the datastream when the tracking mode was active, the weighted centroid position over 12 points (8 ms) was calculated to obtain x and y fluorophore positions for each time section. To avoid building trajectories of noise when no fluorophore was present in the confocal volume, a background threshold was calculated and used to cut multiple trajectories out of the same active tracking mode sequence when the signal was below the background. In a corresponding way as for the EMCCD data analysis, classification of sliding trajectories were done using PCA (see Extended Data Table 5 for classification parameters), were the classifier was optimized to effectively find sliders in flow-channels with DNA but not in negative control channels without DNA (Extended Data Table 4). 1D diffusion constants for sliding trajectories were estimated using the covariance based estimator^25^ from displacements (weighted centroid positions over 160 ms) along the principal component with the largest eigenvalue (the DNA stretching direction).

#### Autocorrelation functions from photon time tagging data

The photon time tagging data was asynchronously recorded through a time-tagged time-resolved (TTTR) data acquisition method. Reconstructing a synchronous time trace of the photon time series is not feasible since this becomes very computational and memory intensive. Instead we used a time-tag-to-correlation algorithm^26^ to directly convert the TTTR data to a fluorescence correlation spectroscopy (FCS) curve. This is done on the summed photon time series from both APDs, effectively recreating the time trace as if there were only one detector. The position trace and autocorrelation of a bead tracking trajectory moving in a circle is shown in Extended Data Figure 5. The autocorrelation data is compensated for the 166 μs off time between laser pulses, meaning that we have only used data when the 500 μs pulse is active. The normalized autocorrelation for each tracking event was calculated from the non normalized autocorrelation *g(τ)* according to

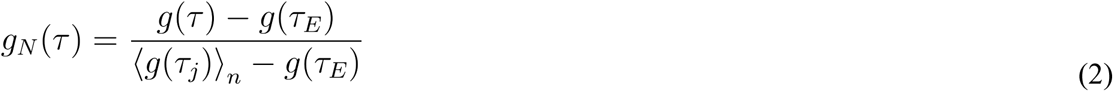

where *τ*_*E*_ is the last time lag of time-scale and the < >_*n*_ operator denotes the average for the first *n* points of the time-scale. For each sliding molecule with high enough quality (see Table Extended Data Table 5 for classification parameters), the normalized autocorrelation function was calculated on four time intervals ranging from 2.5 to 660 µs, with 6 time lags in each interval and with *n* = 2 used for normalization. The average normalized autocorrelation for each protein and time point was then calculated as a 20% trimmed mean of the single molecule data.

#### Vector based dipole point spread function model

The model used for the polarization and orientation dependent emission of a dipole are based on a previous work^18^. Briefly, the polarization intensities in the back focal plane are found by translating the electric far field of the dipole into the back focal plane, and integrating the intensity (square of the electric field) over the numerical aperture *NA* of the objective. After integration we get the polarization intensities

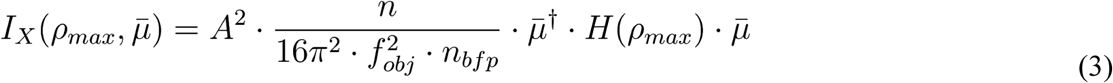

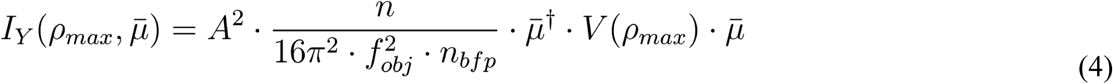

Where *μ* is the dipole orientation, *A* is the amplitude of the dipole moment (dependent on the polarization of the excitation light and the dipole orientation), *n* and *n*_*bfp*_ are refractive indices surrounding the dipole and the back focal plane respectively, *f*_*obj*_ is the focal length of the objective, *ρ*_*max*_ =*NA/n* and *H(ρ*_*max*_) and *V(ρ*_*max*_) are diagonal matrices only dependent on *ρ*_*max*_ (see Supplementary Information for derivation and details). The total intensity detected for this dipole is the sum of these intensities, that is

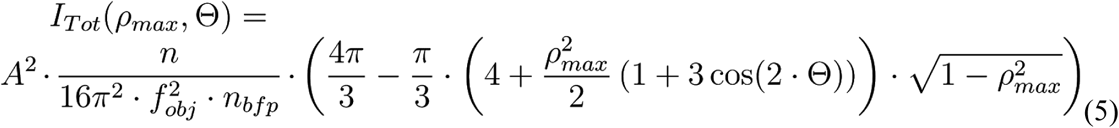

where *Θ* is the polar angle of the dipole orientation (see Fig 1a). The amplitude of the dipole moment was set according to the excitation light of the modelled process (*A* = 1 for isotropic excitation light, *A* = *sinΘ* for excitation light circularly polarized light in the xy-plane).

#### Simulating polarization distributions and autocorrelation functions

The simulated polarization distributions in Figure 2 were generated by calculating polarization intensities according to the vector based dipole point spread function model (Eqs. 3 and 4) while the dipole orientation *μ* was sampled according to the different sliding models (uniform, linear or rotation coupled sliding) and experiment coordinate systems (polarization axis X aligned with average DNA direction or shifted 45° from it). The total number of photons per time step was modeled as a poissonian process with an average photon count rate according to the experimentally measured averages for LacI-BR (see Supplementary Information for details).

The relationship between the autocorrelation function *g(τ)* and the rotational diffusion constant *D*_*r*_ for rotation coupled sliding was found by simulating fluorescence intensity traces emitted from dipoles rotating diffusionaly around the DNA-axis. Brownian dynamics simulations of the rotation of the fluorophore were performed for both 1D (see Supplementary Information) and 3D^27^ rotational diffusion, while the fluorescence intensity was calculated from the vector-based point-spread function model (Eq. 5) at every time-step. After calculating the autocorrelation function from these intensity traces we found that the simulated autocorrelations for 1D rotational diffusion could be described accurately with the model (Extended Data Fig. 7)

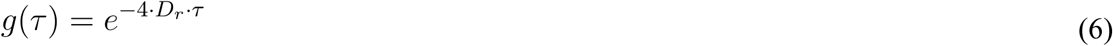

The simulation scheme was validated using the 3D rotational diffusion simulation, where the resulting autocorrelation, as expected, could be described accurately by an exponential with exponent −6*D*_*r*_*τ* (Extended Data Fig. 7).

#### Model fitting and pitch estimation

In rotation coupled sliding, translational movement of the particle is coupled to rotation around the DNA. The rotational diffusion constant *D*_*r*_ and translational diffusion constant *D* are then related to each other according to

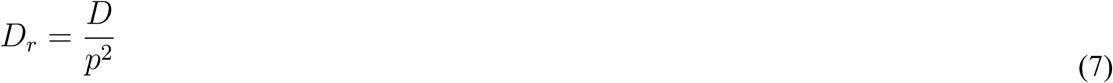

Where *p* is the pitch of the rotational sliding, that is how long the particle translates per angle it rotates around the DNA. To minimize contributions from the background, that is all contributions to the autocorrelation function that are independent of rotational sliding of the molecule, model fitting was done on the difference in autocorrelation between two constructs of LacI (LacI-MBP-BR and LacI-BR) labeled with the same fluorophore but sliding with different rates. Fitting was done on an expression where normalization of the autocorrelation has been done in a corresponding way as for the experimental data

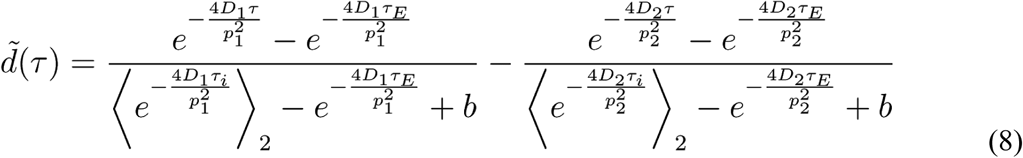

The free parameters of the model are the pitches *p*_*1*_ and *p*_*2*_ of the two proteins and the background parameter *b*, which are fitted by minimizing the square deviation between the experimental difference in autocorrelation and the theoretical difference according to equation 8, in all time regimes simultaneously (see Supplementary Information for details). Translational diffusion constants *D*_*1*_ and *D*_*2*_ were taken as the experimentally measured averages from the tracking data. Rotational pitches have been converted into the unit of bp per full revolution around the DNA in the main-text.

### Molecular dynamics simulations

The structure of the N-terminal domain of LacI bound non-specifically to a 18 nt long dsDNA^7^ was labeled *in silico* by introduction of cysteins and fluorophores at the appropriate positions for LacI-BR and LacI-Cy3 respectively. 100 ns molecular dynamics simulations were then performed using Desmond^28^ version 3.8 and the OPLS-2005 force field (see Supplementary information for details). A burn in time of 30 ns was used to equilibrate the systems before analysis of fluorophore rigidity, so that only the last 70 ns of the simulations were used for analysis. The direction of the fluorophore backbone at each 3 ps snap-shot was calculated as the vector going between the two nitrogen atoms for Cy3, and as the vector going between the two nitrogen atoms closest to the aromatic ring for rhodamine. The spherical coordinates *Θ* and *Φ* of the fluorophore backbone are reported for the coordinate system depicted in Fig. 1a of the main-text, were the x-axis is defined along the DNA direction (calculated from the difference of the DNA end positions) and the z-axis is defined as the vector orthogonal to the x-axis going through the center of mass of the protein.

